## Supplemental Figure for "Subtilase SBT5.2 inactivates flagellin immunogenicity in the plant apoplast"

**Supplemental Figures S1-S10**

Buscaill et al., *Subtilase SBT5.2 inactivates flagellin immunogenicity in the plant apoplast*

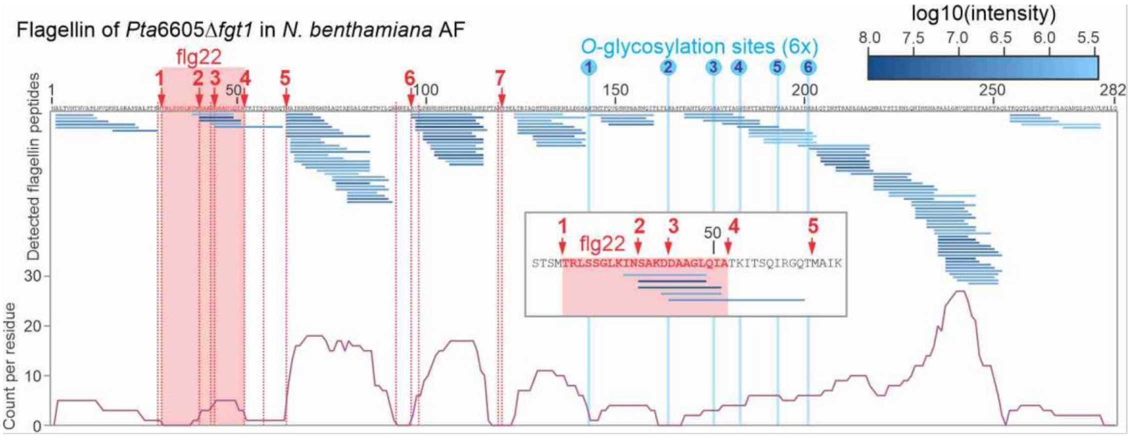


**Fig. S1** Peptide coverage of nonglycosylated flagellin incubated in AF of *N. benthamiana.*

10 µg/ml of purified flagellin of *Pta*6605Δ*fgt1* was incubated for 60 minutes with apoplastic fluids isolated from agroinfiltrated *N. benthamiana* leaves expressing the empty vector (EV). Proteins were precipitated with 80% acetone and the peptide fraction (supernatant) was analysed by mass spectrometry. Flagellin-derived peptides were aligned to the flagellin protein sequence. Highlighted are the flg22 sequence, putative cleavage sites (red lines) and six *O*-glycosylation sites (blue lines). The number of times each residue was detected in the peptides is indicated with the purple graph, showing the peptides belong to seven clusters. Inset: region containing flg22 and cleavage sites 1-5 with the corresponding detected peptides.

| Name | NbDE* | Mock control (5dpi) | | Agroinfiltrated (5pi) | |
|---|---|---|---|---|---|
| | | mean | SD | mean | SD |
| *Nb*SBT5.2a | NbD038072 | 0 | 0 | 0.093467031 | 0.042122779 |
| *Nb*SBT5.2b | NbD021558 | 9.744890966 | 2.14370341 | 4.553243162 | 0.921318851 |
| *Nb*SBT5.2c | NbD013006 | 0.618187564 | 0.27390427 | 0.523724281 | 0.303865567 |

**Fig. S2** Transcript levels of *SBT5.2* genes in *N. benthamiana* leaves.

Reads Per Kilobase of transcript per Million mapped reads (RPKM) values of *Nb*SBT5.2a, *Nb*SBT5.2b, and *Nb*SBT5.2c in mock or agroinfiltrated plants[35]. *, NbDE annotation[17].

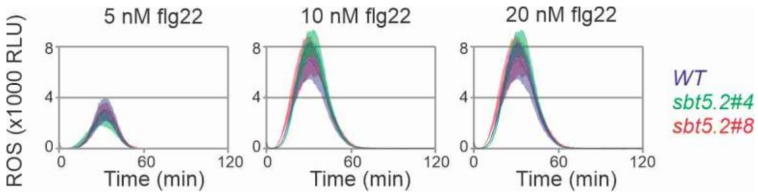


**Fig. S3** No altered response to flg22 in *sbt5.2* mutants.

Leaf discs from 4-week-old *N. benthamiana* WT or *sbt5.2* mutants floating on luminol-HRP were treated with 5, 10 or 20 nM flg22 and ROS burst was monitored using a plate reader. Error shades represent SE of n=6 biological replicates.

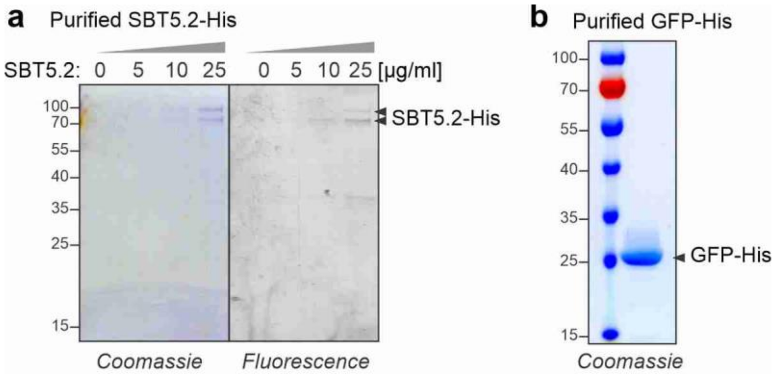


**Fig. S4** Purified SBT5.2a-His an GFP-His.

His-tagged SBT5.2a and secreted GFP were transiently expressed in *N. benthamiana* by agroinfiltration and isolated from AF on Ni-NTA columns. **a.** Different SBT5.2-His concentrations were labelled with 0.5 µM FP-TAMRA for one hour and samples were separated on SDS-PAGE and scanned for in-gel fluorescence. **b.** Purified GFP-His was separated on SDS-PAGE and stained with Coomassie.

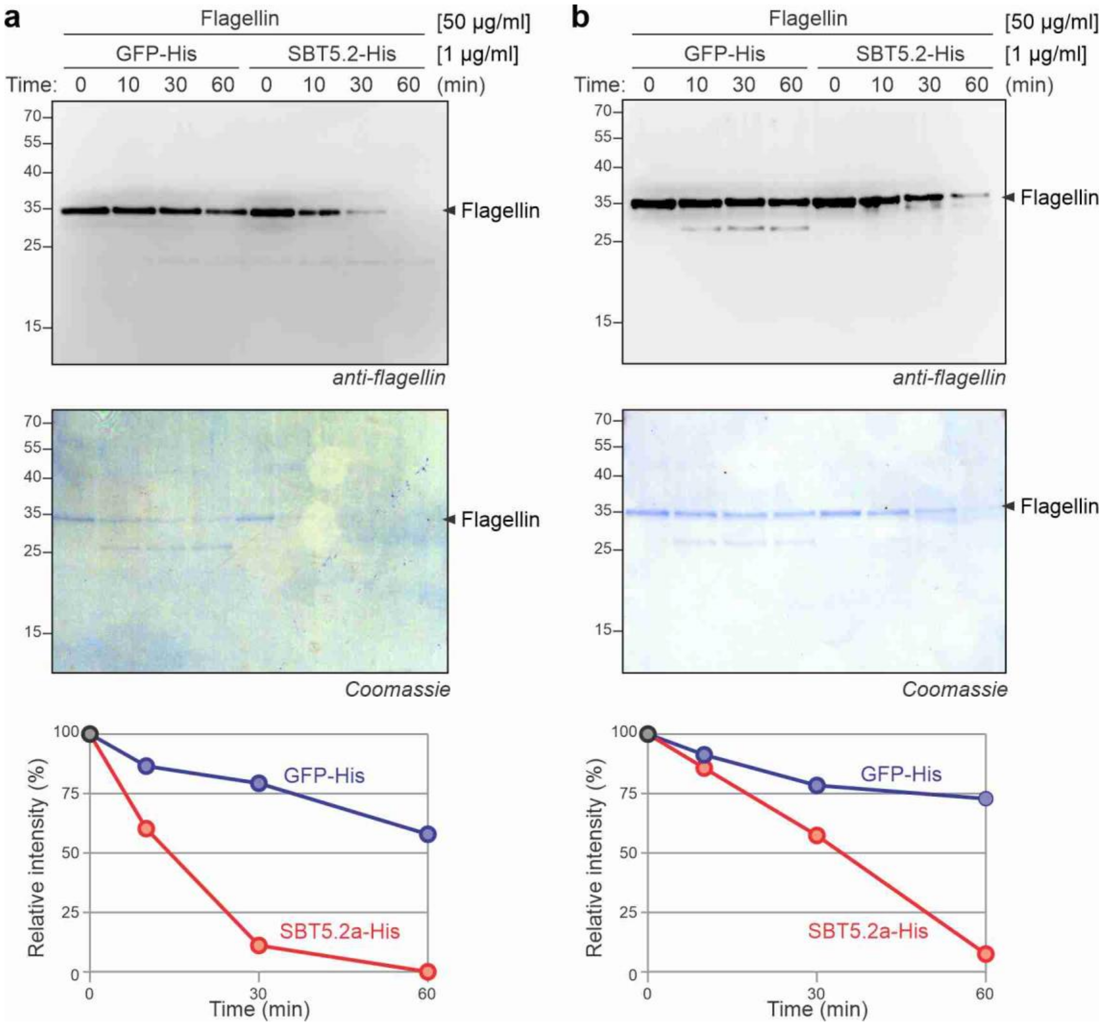


**Fig. S5** Purified SBT5.2a degrades flagellin protein.

Purified SBT5.2a-His processes flagellin. Purified flagellin (50 µg/ml) was incubated with of purified SBT5.2a-His or GFP-His (1 µg/ml) for 0, 10, 30 or 60 minutes and samples were separated on SDS-PAGE and stained with Coomassie or analysed by western blotting using anti-flagellin antibodies. Shown are two replicate experiments, with western blot (top), Coomassie-stained gel (middle) and quantification of western blot signals relative to t=0 (bottom).

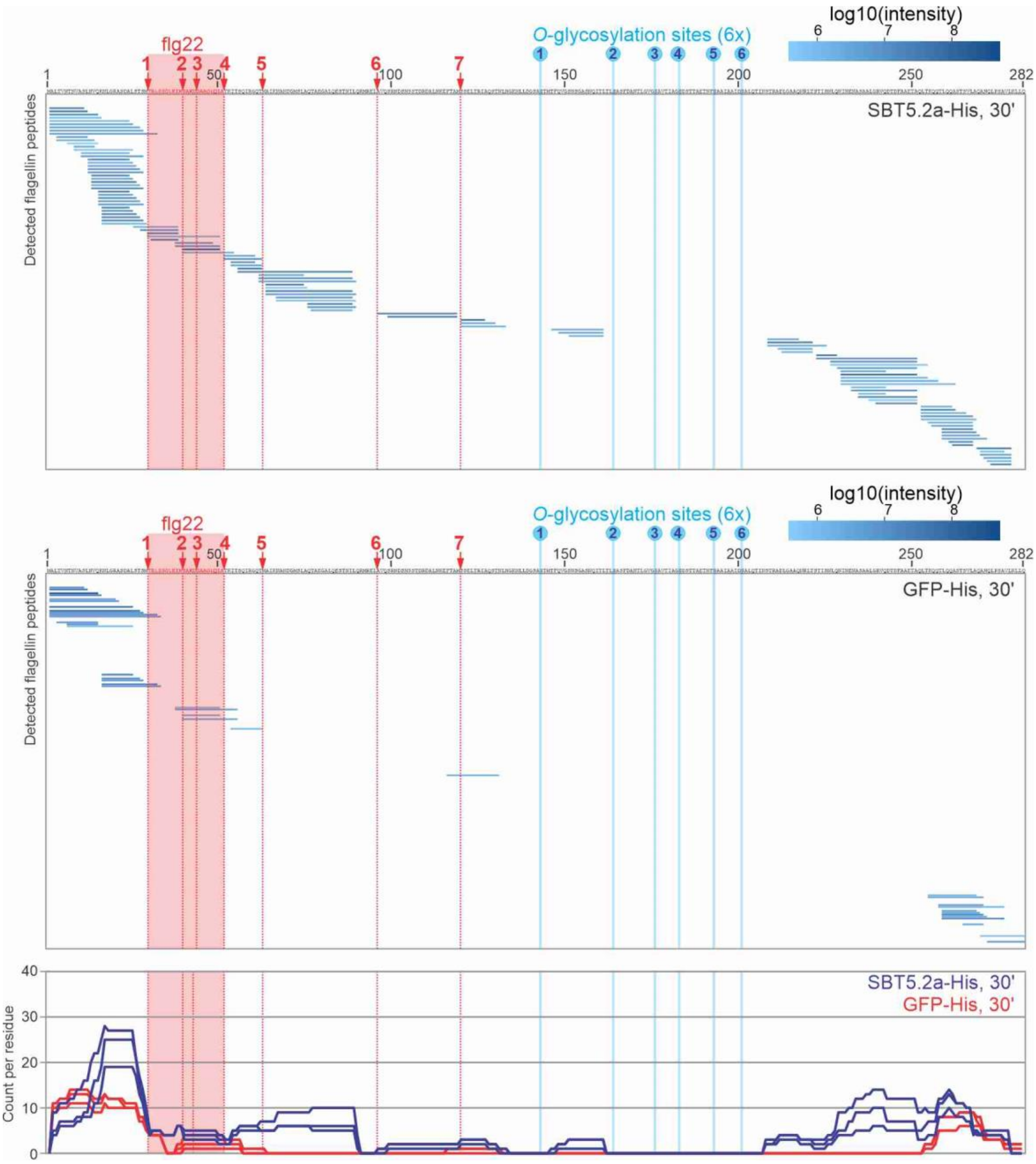


**Fig. S6** Purified SBT5.2a-His cleaves flagellin in the flg22 epitope.

Flagellin was incubated with purified SBT5.2-His of GFP-His for 30 minutes and the released peptides were analysed by LC-MS/MS. Shown is the mean of n=3 replicates. The bottom graph summarises the count for each residue detected in the three replicates.

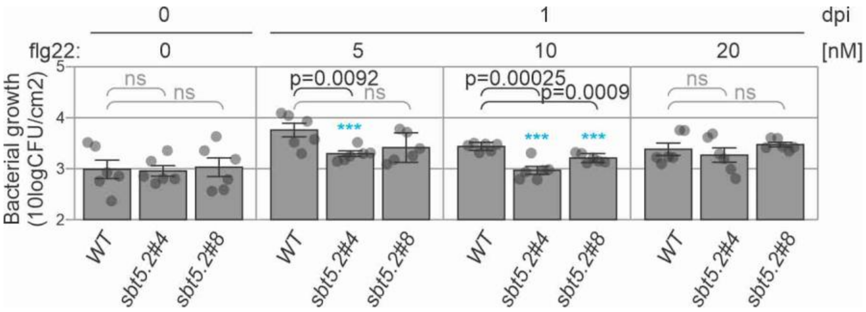


**Fig. S7** Immune priming by low flg22 concentrations increases in *sbt5.2* mutant plants. Leaves of 4-week-old WT and *sbt5.2* mutant plants were infiltrated with 5, 10 or 20 nM flg22 or water. After 24 hours incubation, the leaves were infiltrated with $1x10^5$ bacteria/ml *Pta*6605. Colony forming units (CFUs) were determined one day post infection (dpi). Error bars represent SE of n=6 replicates.

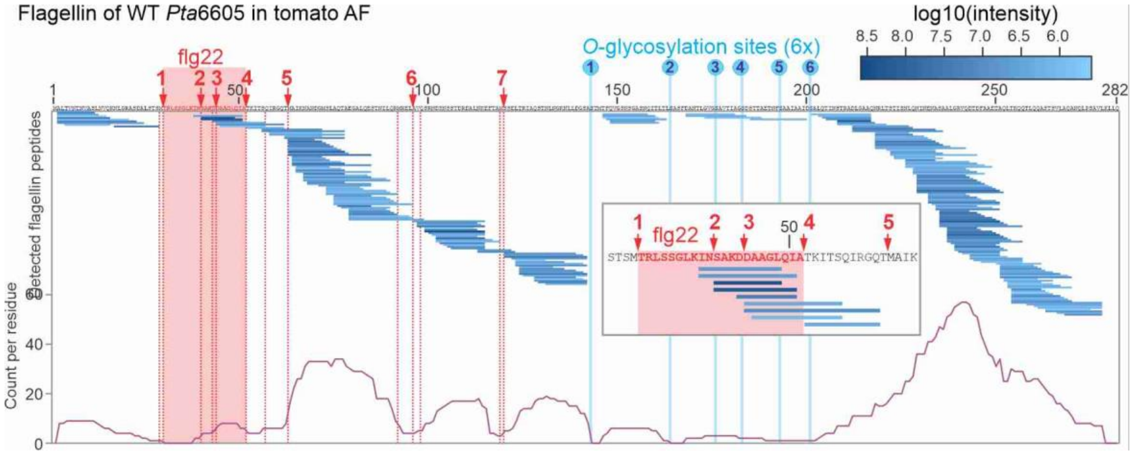


**Fig. S8** Peptide coverage of flagellin incubated in AF of tomato.

10 µg/ml of purified flagellin of *Pta*6605 was incubated for 60 min with apoplastic fluids isolated from tomato (Money Maker Cf0). Proteins were precipitated with 80% acetone and the peptide fraction (supernatant) was analysed mass spectrometry. Flagellin-derived peptides were aligned with the flagellin protein sequence. Highlighted are the flg22 sequence, putative cleavage sites (red lines) and six *O*-glycosylation sites (blue lines). The number of times each residue was detected in the peptides is indicated with the purple graph, showing the peptides belong to seven clusters. Inset: region containing flg22 and cleavage sites 1-5 with the corresponding detected peptides.

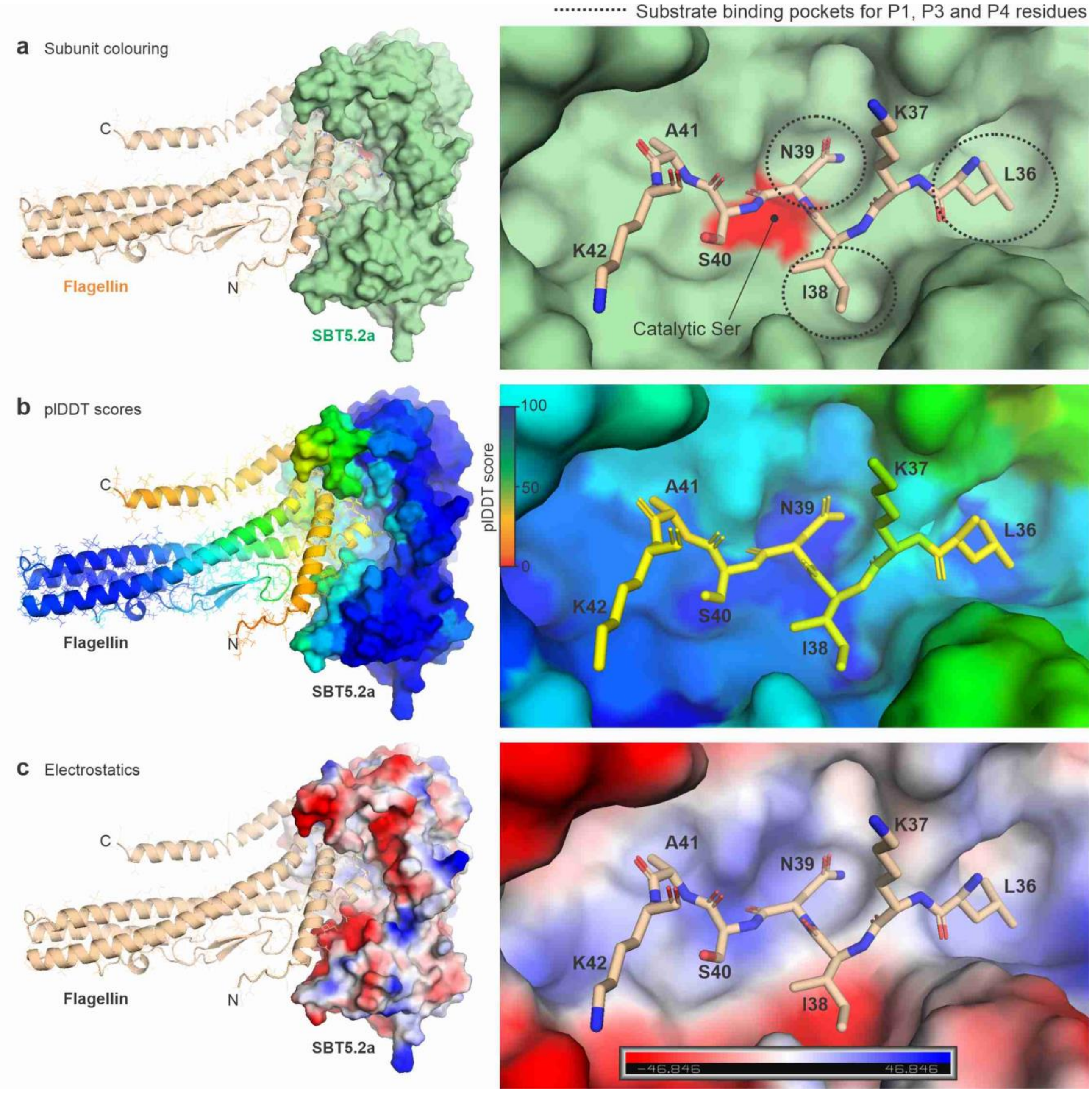


**Fig. S9** Structural model of flagellin interacting with SBT5.2a.

**a.** Colouring for subunit annotation. **b.** Colouring by plDDT scores of both proteins. **c.** Colouring by vacuum electrostatics of SBT5.2a. Left: overview of the complexes; right: zoom in on the LKINSAK peptide of flagellin bound to the substrate binding groove of SBT5.2a. AlphaFold Multimer was used to predict complexes between the catalytic domain of SBT5.2a and flagellin. The best ranking complex had a moderate score (ipTM+pTM=0.51) and relatively low plDDT scores in interface region of flagellin, but the plDDT scores of SBT5.2a are high. Strikingly, the flagellin monomer is not rod-like but has bend on the flg22 hinge, exposing the preferred cleavage site (LKIN'SAK) to the substrate binding groove of SBT5.2a, with the cleavable bond in close proximity to the catalytic Ser (red in A), assigning N39, I38 and L36 to the P1, P2 and P4 positions, respectively, consistent with the observed cleavage.

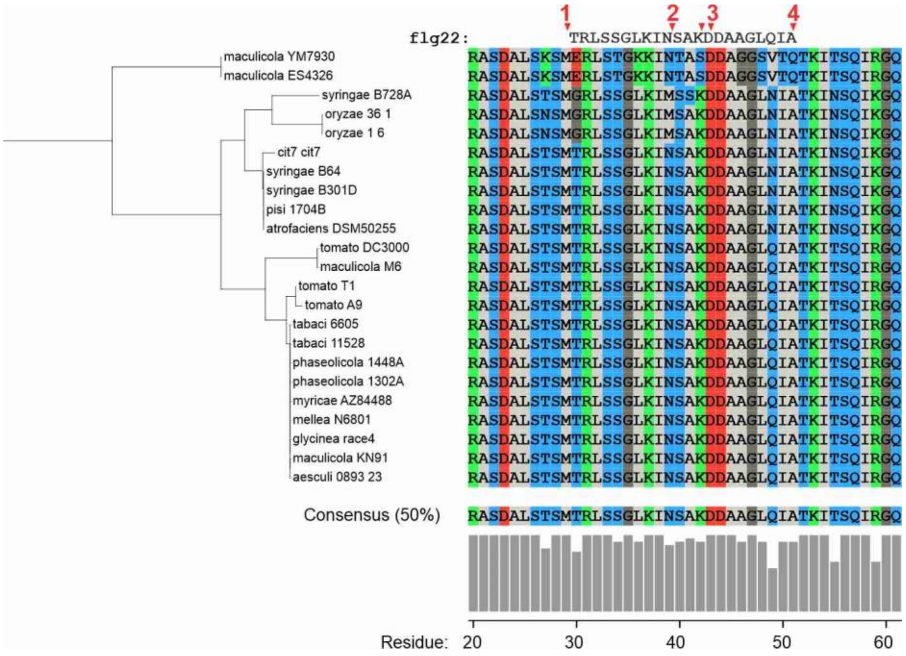


**Fig. S10** Processing sites 1-4 are highly conserved in the flg22 epitope.

Phylogenetic tree of flagellin proteins from *Pseudomonas syringae* pathovars. Clustal Omega was used for amino acid sequence alignment and neighbour-joining tree construction. The tree was visualised using iTOL and displayed with midpoint rooting.
