## Supplementary material for "Subtilase SBT5.2 inactivates flagellin immunogenicity in the plant apoplast": Table S5

### Supplemental Tables S5

Buscaill et al., *Subtilase SBT5.2 inactivates flagellin immunogenicity in the plant apoplast*

**Table S5** Used (quenched) peptides

| **Peptide** | **Sequence** | **Reference** |
|---|---|---|
| QP1 | DABCYL-STSMTRLS-(G)EDANS | This study |
| QP2 | DABCYL-LKINSAKD-(G)EDANS | This study |
| QP3 | DABCYL-SAKDDAAG-(G)EDANS | This study |
| QP4 | DABCYL-LQIATKITS-(G)EDANS | This study |
| QP5 | DABCYL-RGQTMAIK-(G)EDANS | This study |
| flg22 | TRLSSGLKINSAKDDAAGLQIA | This study |
