## Supplementary material for "Subtilase SBT5.2 inactivates flagellin immunogenicity in the plant apoplast": Table S6

**Supplemental Table S6**

Buscaill et al., *Subtilase SBT5.2 inactivates flagellin immunogenicity in the plant apoplast*

**Table S6** Plasmids used in this work.

| **Plasmid** | **Description** | **Reference** |
|---|---|---|
| pFGH48 | 35S::Epi1 | Grosse-Holz et al., 2018[16] |
| pFGH54 | 35S::*Sl*CYS8 | Grosse-Holz et al., 2018[16] |
| pFGH47 | 35S::*Hs*TIMP | Grosse-Holz et al., 2018[16] |
| pPB097 | *35S::SBT5.2a-HIS* | Chen et al., 2024[22] |
| pNS205 | *35S::SP-GFP-His* | This study |
| pJK187 | binary vector p35S::LacZ::t35S | Homma et al., 2023[41] |
| pJK037 | pL2M-TRV2 | Morimoto et al., 2022 |
| pPB039 | *TRV2::SBT5.2* | Beritza et al., 2024[18] |
| pPB058 | *TRV2::SBT1.7a* | Beritza et al., 2024[18] |
| pPB059 | *TRV2::SBT1.7c* | Beritza et al., 2024[18] |
| pPB065 | *TRV2::SBT1.9a* | Beritza et al., 2024[18] |
