## Supplementary material for "Subtilase SBT5.2 inactivates flagellin immunogenicity in the plant apoplast": Table S7

**Supplemental Table S7**

Buscaill et al., *Subtilase SBT5.2 inactivates flagellin immunogenicity in the plant apoplast*

**Table S7** Oligonucleotides used in this work

| Use | Sequence (5'-3') |
|---|---|
| PR1a-GFP-His | TTGAAGACTCAATGGGTTTCGTGCTGTTCTCTCAGCTGCCTTCTTTCCTTCTTGTGTCTACCCTTCTGCTGTTCCTGGTGATCTCTCATTCTTGCAGGGCTCAGAACTCTGGTCATCATCATCACCATCACGGCAGCATGAGGAAGGGTGAAGAGTTGTTCACTGGTGTGGTGCCTATTCTGGTTGAGCTTGATGGGGATGTGAACGGCCATAAGTTCAGCGTTAGAGGTGAAGGTGAGGGTGATGCTACCAACGGTAAGCTGACCCTTAAGTTCATCTGTACCACCGGAAAGTTGCCTGTGCCTTGGCCTACTCTTGTGACCACTCTTACTTACGGTGTGCAGTGCTTCGCTAGGTATCCTGATCATATGAAGCAGCACGACTTCTTCAAGAGCGCTATGCCTGAGGGTTACGTGCAAGAGAGGACCATCAGCTTCAAGGATGATGGCACTTACAAGACCCGTGCCGAGGTTAAGTTCGAGGGTGATACTCTGGTGAACCGGATTGAGCTGAAGGGCATCGATTTCAAAGAGGACGGTAACATCCTGGGCCACAAGCTCGAGTACAACTTCAACTCTCACAACGTGTACATCACCGCCGACAAGCAGAAGAACGGTATCAAGGCCAACTTCAAGATCCGGCACAACGTTGAGGATGGTTCTGTGCAGCTTGCTGATCACTACCAGCAGAACACCCCTATTGGTGATGGACCTGTGCTTCTGCCTGATAACCACTACCTTTCTACCCAGAGCGTGCTGAGCAAGGATCCTAATGAGAAGAGGGATCACATGGTGCTGCTTGAGTTCGTTACCGCTGCTGGTATTACCCACGGTATGGATGAGCTGTACAAGGGCTCTGCTTGGTCACATCCTCAGTTCGAGAAGTAGGCTTGAGTCTTCAA |
