## Supplementary material for "Subtilase SBT5.2 inactivates flagellin immunogenicity in the plant apoplast": Data S1

### Overview

| ACE project | Title |
|---|---|
| ACE_0383 | Comparison of mass spectra among flagellin protein samples |
| ACE_0481 | Hydrolytic release of flagellin peptides. |
| ACE_0935 | Released peptide from flagellin cleavage by SBT5.2a |

### ACE_0383

#### File legend

| ACE ID | Organism | Organ/ cell line | Treatment/ experimental setup |
|---|---|---|---|
| ACE_0383_PB10 | Pseudomonas | flagella | 25ul of apoplastic fluids of N. benthamiana transiently expressing P19 mixed with the flagellin proteins isolated from Pseudomonas Pta6605 WT (at 10ng/ul) |
| ACE_0383_PB11 | Pseudomonas | flagella | 25ul of apoplastic fluids of N. benthamiana transiently expressing P19 mixed with the flagellin proteins isolated from Pseudomonas Pta6605 WT (at 10ng/ul) |
| ACE_0383_PB12 | Pseudomonas | flagella | 25ul of apoplastic fluids of N. benthamiana transiently expressing P19 mixed with the flagellin proteins isolated from Pseudomonas Pta6605 WT (at 10ng/ul) |

#### LC_Settings

| | |
|---|---|
| MS device | Orbitrap Elite |
| LC device | Thermo Easy-nLC 1000 |
| ion source | Thermo Nanospray Flex |
| **Analytical column** | Self-packed fused silica capillary with integrated pico frit emitter; New Objectives PF360-75-15-N-5 |
| column diameter | Length ($L_C$) = 35 cm; ID = 75µm; OD = 360 µm; emitter 15 µm |
| stationary phase | Reprosil-Pur 120 C18-AQ, Dr. Maisch GmbH |
| particle diameter ($d_p$) | 1.9 µm |
| Pore size | 120 Å |
| Column ID | AC61 |
| Column oven | Sonation column oven PRSO-V1 |
| Column oven temp. | 45°C |
| **solvents** | A: 0.1% FA in UPLC water<br>B: 0.1% FA in 80% UPLC ACN |
| gradient | 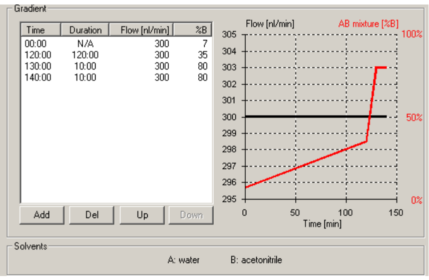 |

#### MS_Settings

| Project | MS | general | MS1 | MS2 | MS2 | MS3 | Comments; special settings |
|---|---|---|---|---|---|---|---|
| ACE_0383 | Elite | Tune v2.7.0.1112 SP2<br>Xcalibur: v3.0.63<br>Gradient: 140 min | Analyzer: FT<br>Res.: 60000<br>SR: 300 - 1700<br>AGC: $3\times10^6$<br>AcT: 50 ms<br>RF: n/a<br>SF: --<br>DDM: NS/15 | Analyzer: IT<br>Res./ScR: -/rapid<br>SR: Auto<br>AGC: $1\times10^4$<br>AcT: 50 ms<br>CS: > +2<br>IsM: IT<br>IsW: 2.0<br>Frag.: CID<br>NCE: 35 | | | classic orbitrap experiment: MS1 in Orbitrap at high resolution and data dependent MS2 in Iontrap at rapid scan rate. Dynamic exclusion enabled (exclude after n times=1; Exclusion duration (s)= 120; mass tolerance= ± 10 ppm)<br><br>capillary temp.: 200 °C |

Note: **FT**= Fourier Transform (Orbitrap); **IT**= Iontrap; **Q**= Quadrupol; **Res.**= max. Resolution at 200 m/z (Lumos) or 400 m/z (Elite) [FWHM (full width at half maximum)]; **ScR**= scan rate for measurements in the IT; **SR**= scan range [m/z]; **AGC**= automatic gain control, max number of acquired ions per measurement; **AcT**= max. Ion acquisition time [ms]; **CS**= charge states used for fragmentation; **IsM**= Isolation mode (Q or IT), MS2 isolation and further is only done in IT; **IsW**= Isolation window [m/z], value followed by scan mode the isolation is based on (MS1, MS2 …) **Frag.**= Fragmentation method; **HCD**= Higher-energy collisional dissociation; **CID**= Collision-induced dissociation; **ETD**= Electron-transfer dissociation**; EThcD=** Electron-Transfer/Higher-Energy Collision Dissociation; **sHCD**= stepped HCD**; NCE**= normalized collision energy; **cycles**: number of MSn recorded or max cycle time; RF= RF Lens [%]; **SF**= Source Fragmentation [V]; **DDM**: Data dependent Mode (cycle time in seconds, CT/[s] or number of scans, NS); **NS**= Number of data dependent scans

#### Search Settings

| | |
|---|---|
| Program & version | MaxQuant v1.5.5.30. |
| Search engine | Andromeda |
| settings | Basically default; LFQ and MBR were turned on |
| Static modification | none |
| Digestion mode | unspecific |
| Dynamic modification | Acetyl (N-term); Oxidation (M) |
| Modification included in quantification | Oxidation (M) |
| Databases | 1. Contaminants<br>2. p19_vector_proteins.fasta<br>3. ACE_0383_SOI_v02.fasta |

### ACE_0481

#### File legend

| ACE ID | Organism | Organ/ cell line | Treatment/ experimental setup |
|---|---|---|---|
| ACE_0481_PB01 | *Pseudomonas* | flagella | 25 µl of AF from *N. benthamiana* overexpressing P19+**EV** and mixed with the flagellin proteins isolated from Pseudomonas ***Pta*6605Δ*fgt1*** (at 10ng/µl) |
| ACE_0481_PB05 | *Pseudomonas* | flagella | 25 µl of AF from Tomato incubated with **DMSO** and mixed with the flagellin proteins isolated from Pseudomonas ***Pta*6605** (at 10ng/µl) |

#### LC_Settings

| | |
|---|---|
| MS device | Thermo Orbitrap Fusion Lumos |
| LC device | Thermo Easy-nLC 1200 |
| ion source | Thermo Nanospray Flex |
| **Analytical column** | Self-packed fused silica capillary with integrated pico frit emitter; New Objectives PF360-75-15-N-5 |
| column diameter | Length ($L_C$) = 30 cm; ID = 75µm; OD = 360 µm; emitter 15 µm |
| stationary phase | Reprosil-Pur 120 C18-AQ, Dr. Maisch GmbH |
| particle diameter ($d_p$) | 1.9 µm |
| Pore size | 120 Å |
| Column ID | Ac82 |
| Column oven | Sonation column oven PRSO-V2 |
| Column oven temp. | 50°C |
| **solvents** | A: 0.1% FA in UPLC water<br>B: 0.1% FA in 80% UPLC ACN |
| gradient | 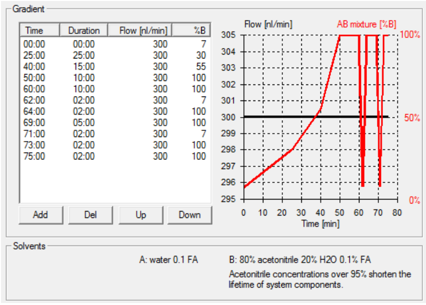 |

#### MS_Settings

| Project | MS | general | MS1 | MS2 | MS2 | MS3 | Comments; special settings |
|---|---|---|---|---|---|---|---|
| ACE_0481 | Lumos | Tune v3.5.3881.18<br>Xcalibur v4.5.445.18<br>Gradient: 75 min | Analyzer: FT<br>Res.: 120000<br>SR: 375 - 1500<br>AGC: Standard<br>AGC abs.: 400000<br>AcT: 50 ms<br>RF: 30<br>SF: --<br>DDM: CT/3sec | Analyzer: IT<br>Res./ScR: -/rapid<br>SR: Auto<br>AGC: 100%<br>AGC abs.: 10000<br>AcT: 35 ms<br>CS: +2 to +7<br>IsM: Q<br>IsW: 1.2<br>Frag.: CID<br>NCE: 35 | | | classic orbitrap experiment: MS1 in Orbitrap at high resolution and data dependent MS2 in iontrap. Dynamic exclusion enabled (exclude after n times=1; Exclusion duration (s)= 60; mass tolerance= ± 10ppm)<br><br>Intensity Threshold: 5000<br>Ion transfer Tube Temp: 275 °C<br>Ion Source Voltage: 2500 V |

Note: **FT**= Fourier Transform (Orbitrap); **IT**= Iontrap; **Q**= Quadrupol; **Res.**= max. Resolution at 200 m/z (Lumos) or 400 m/z (Elite) [FWHM (full width at half maximum)]; **ScR**= scan rate for measurements in the IT; **SR**= scan range [m/z]; **AGC**= automatic gain control, max number of acquired ions per measurement; **AcT**= max. Ion acquisition time [ms]; **CS**= charge states used for fragmentation; **IsM**= Isolation mode (Q or IT), MS2 isolation and further is only done in IT; **IsW**= Isolation window [m/z], value followed by scan mode the isolation is based on (MS1, MS2 …) **Frag.**= Fragmentation method; **HCD**= Higher-energy collisional dissociation; **CID**= Collision-induced dissociation; **ETD**= Electron-transfer dissociation**; EThcD=** Electron-Transfer/Higher-Energy Collision Dissociation; **sHCD**= stepped HCD**; NCE**= normalized collision energy; **cycles**: number of MSn recorded or max cycle time; RF= RF Lens [%]; **SF**= Source Fragmentation [V]; **DDM**: Data dependent Mode (cycle time in seconds, CT/[s] or number of scans, NS); **NS**= Number of data dependent scans

#### Search Setting

| | |
|---|---|
| Program & version | MaxQuant v1.6.3.4 |
| Search engine | Andromeda |
| settings | Basically default; LFQ and MBR were turned on |
| Static modification | none |
| Digestion mode | unspecific |
| Dynamic modification | Acetyl (N-term); Oxidation (M) |
| Modification included in quantification | Oxidation (M) |
| Databases | 1. Contaminants<br>2. ACE_0481_SOI_v02.fasta<br>3. p19_vector_proteins.fasta<br>4. 12864_2019_6058_MOESM10_ESM.fasta (*N. benthamiana*))<br>5. UP000000813_176299.fasta (*A. tumefaciens strain C58*) |

### ACE_0935

#### File legend

| ACE ID | Organism | Organ/ cell line | Treatment/ experimental setup |
|---|---|---|---|
| ACE_0935_PB04 | *N. benth;P. syringae* | | Flagellin +GFP 30 min - replicate 1 |
| ACE_0935_PB05 | *N. benth;P. syringae* | | Flagellin +GFP 30 min - replicate 2 |
| ACE_0935_PB06 | *N. benth;P. syringae* | | Flagellin +GFP 30 min - replicate 3 |
| ACE_0935_PB13 | *N. benth;P. syringae* | | Flagellin +SBT5.2a 30 min - replicate 1 |
| ACE_0935_PB14 | *N. benth;P. syringae* | | Flagellin +SBT5.2a 30 min - replicate 2 |
| ACE_0935_PB15 | *N. benth;P. syringae* | | Flagellin +SBT5.2a 30 min - replicate 3 |

Note: HIS-tagged GFP and SBT5.2a were expressed in N. benthamiana leaves and purified from leaves AF. Native flagellin proteins were isolated from Pseudomonas syringae pv. tabaci strain 6605.

#### LC_Settings

| | |
|---|---|
| MS device | Orbitrap Fusion Lumos |
| LC device | Thermo Vanquish Neo |
| ion source | Thermo Nanospray Flex |
| **Analytical column** | Self-packed fused silica capillary with an integrated sintered frit; CoAnn Technologies ICT36007515F-50-5 |
| column diameter | Length ($L_C$) = 28 cm; ID = 75µm; OD = 360 µm; emitter 15 µm |
| stationary phase | Phenomenex Kinetex C18-XB core shell |
| particle diameter ($d_p$) | 1.7 µm |
| Pore size | 120 Å |
| Column ID | AC157/AC158 |
| Column oven | Sonation column oven PRSO-V2 |
| Column oven temp. | 50°C |
| **solvents** | A: 0.2% FA, 2% ACN, 98% $H_2O$<br>B: 0.2% FA, 80% ACN, 20 % $H_2O$ |
| gradient | 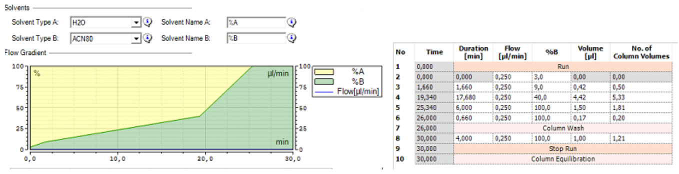 |

| No | Time | Duration [min] | Flow [µl/min] | %B | Volume [µl] | No. of Column Volumes |
|---|---|---|---|---|---|---|
| 1 | 0,000 | Run | | | | |
| 2 | 0,000 | 0,000 | 0,250 | 3,0 | 0,00 | 0,00 |
| 3 | 1,660 | 1,660 | 0,250 | 9,0 | 0,42 | 0,50 |
| 4 | 19,340 | 17,680 | 0,250 | 40,0 | 4,42 | 5,33 |
| 5 | 25,340 | 6,000 | 0,250 | 100,0 | 1,50 | 1,81 |
| 6 | 26,000 | 0,660 | 0,250 | 100,0 | 0,17 | 0,20 |
| 7 | 26,000 | Column Wash | | | | |
| 8 | 30,000 | 4,000 | 0,250 | 100,0 | 1,00 | 1,21 |
| 9 | 30,000 | Stop Run | | | | |
| 10 | 30,000 | Column Equilibration | | | | |

#### MS_Settings

| Project | MS | general | MS1 | MS2 | MS2 | MS3 | Comments; special settings |
|---|---|---|---|---|---|---|---|
| ACE_0935 | Lumos | Tune v4.1.4244<br>Xcalibur v4.7.69.37<br>SII: 1.7.0.468<br>Gradient: 30 min | Analyzer: FT<br>Res.: 120000<br>SR: 375 - 1500<br>AGC: Standard<br>AGC abs.: 400000<br>AcT: 50 ms<br>RF: 30<br>SF: --<br>DDM: CT/3sec | Analyzer: IT<br>Res./ScR: -/rapid<br>SR: Auto<br>AGC: 300%<br>AGC abs.: 30000<br>AcT: 35 ms<br>CS: +2 to +7<br>IsM: Q<br>IsW: 1.6<br>Frag.: HCD<br>NCE: 32 | | | classic orbitrap experiment: MS1 in Orbitrap at high resolution and data dependent MS2 also in Orbitrap high resolution. Dynamic exclusion enabled (exclude after n times=1; Exclusion duration (s)= 20; mass tolerance= ± 10ppm)<br><br>Intensity Threshold: 2000<br>Ion transfer Tube Temp: 250 °C<br>Ion Source Voltage: 2500 V |

Note: **FT**= Fourier Transform (Orbitrap); **IT**= Iontrap; **Q**= Quadrupol; **Res.**= max. Resolution at 200 m/z (Lumos) or 400 m/z (Elite) [FWHM (full width at half maximum)]; **ScR**= scan rate for measurements in the IT; **SR**= scan range [m/z]; **AGC**= automatic gain control, max number of acquired ions per measurement; **AcT**= max. Ion acquisition time [ms]; **CS**= charge states used for fragmentation; **IsM**= Isolation mode (Q or IT), MS2 isolation and further is only done in IT; **IsW**= Isolation window [m/z], value followed by scan mode the isolation is based on (MS1, MS2 …) **Frag.**= Fragmentation method; **HCD**= Higher-energy collisional dissociation; **CID**= Collision-induced dissociation; **ETD**= Electron-transfer dissociation**; EThcD=** Electron-Transfer/Higher-Energy Collision Dissociation; **sHCD**= stepped HCD**; NCE**= normalized collision energy; **cycles**: number of MSn recorded or max cycle time; RF= RF Lens [%]; **SF**= Source Fragmentation [V]; **DDM**: Data dependent Mode (cycle time in seconds, CT/[s] or number of scans, NS); **NS**= Number of data dependent scans

#### Search Settings

| | |
|---|---|
| Program & version | MaxQuant v2.5.2.0 |
| Search engine | Andromeda |
| settings | Basically default; LFQ and MBR were turned on |
| Static modification | none |
| Digestion mode | unspecific |
| | Min. peptide length for unspecific search 7<br>Max. peptide length for unspecific search 36 |
| Dynamic modification | Acetyl (N-term); Oxidation (M) |
| Modification included in quantification | Oxidation (M);Acetyl (Protein N-term) |
| Databases | 1. Contaminants<br>2. ACE_0935_SOI_v01.fasta |
