## Supplementary figures and images for "Subtilase SBT5.2 inactivates flagellin immunogenicity in the plant apoplast"

### Data S2

Fig1a

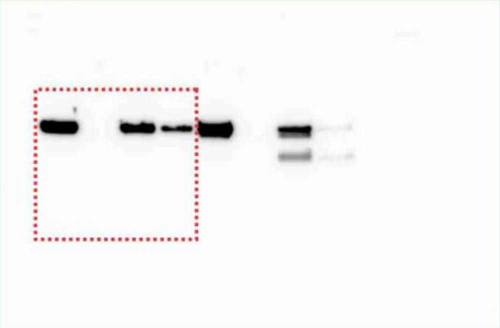

Fig1a

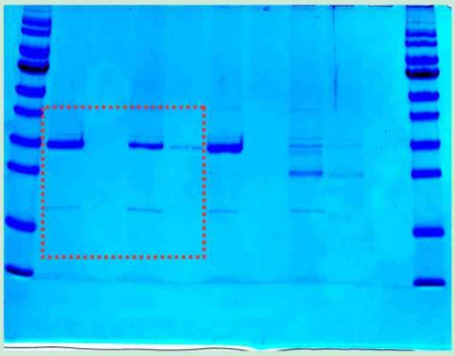

Fig1d

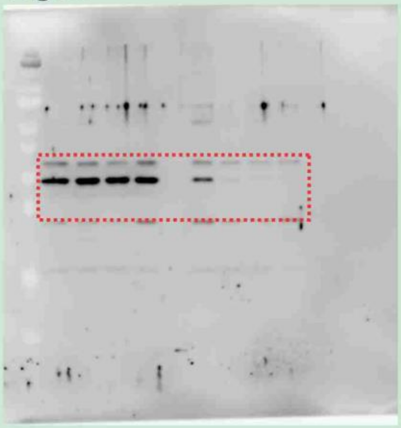

Fig1d

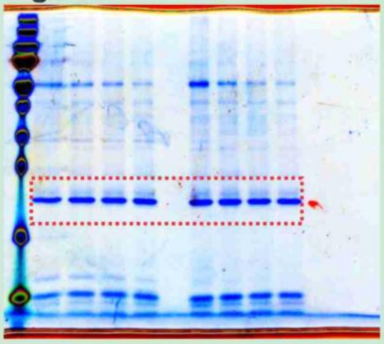

Fig4a

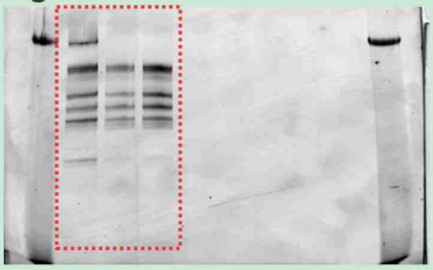

Fig5d

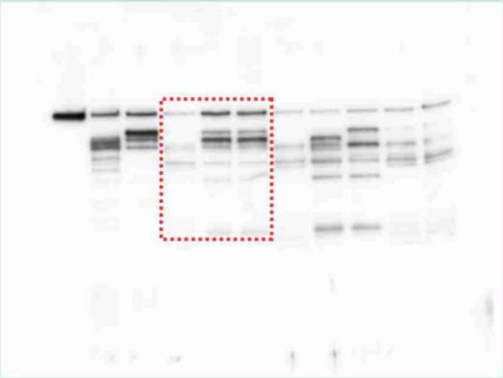

Fig5d

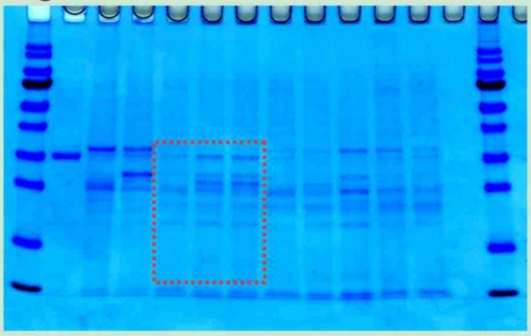

FigS4a

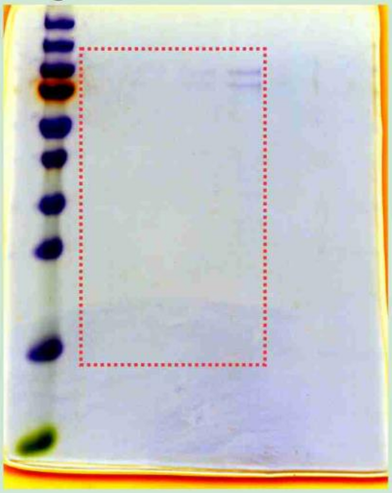

FigS4a

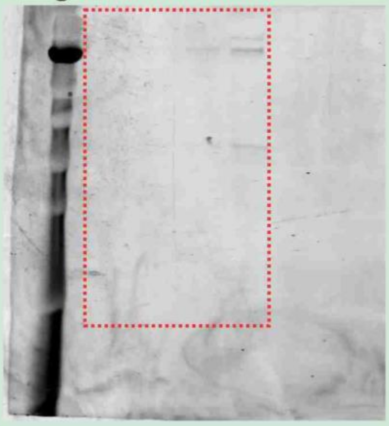

FigS4b

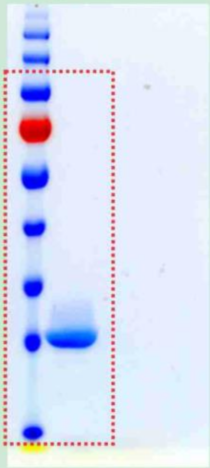

Fig5c/S5

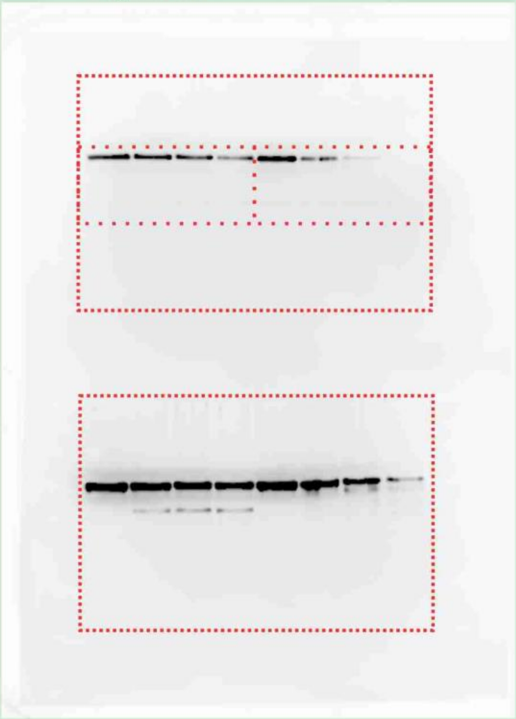

FigS5

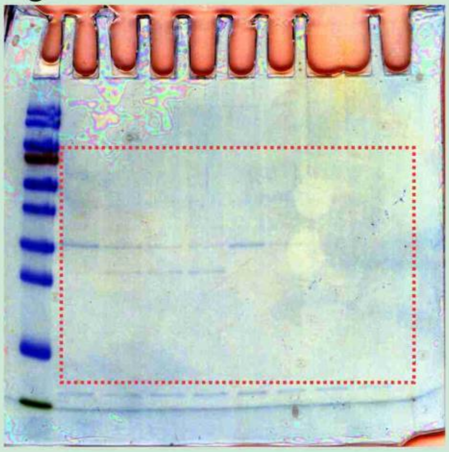

FigS5

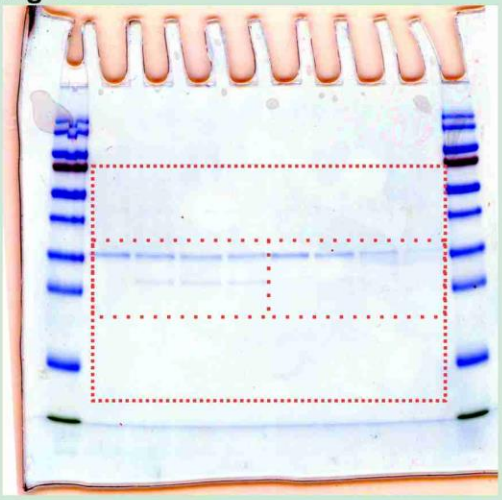
